# Generalizability in White Blood Cells’ Classification Problem

**DOI:** 10.1101/2021.05.12.443717

**Authors:** Sajad Tavakoli, Ali Ghaffari, Zahra Mousavi Kouzehkanan

**Affiliations:** Faculty of Electrical Engineering, K. N. Toosi University of Technology, Tehran, Iran; Nimaad Health Equipment Development Company, Tehran, Iran: www.Nimaadmed.com; Faculty of Mechanical Engineering, K. N. Toosi University of Technology, Tehran, Iran; School of ECE, College of Engineering, University of Tehran, Tehran, Iran

**Keywords:** white blood cells, deep convolutional networks, generalizability, classification, segmentation

## Abstract

Counting and classifying white blood cells (WBCs) in blood samples helps the early diagnosis of the disease. Many works have been done to develop machine learning-based methods to count WBCs. However, most of these works have low generalizability, and their accuracy decreases sharply as the dataset changes. In this paper, a new method is presented that helps to increase the generalization power. In this method, first, the WBC’s nucleus is segmented, and then its convex hull is obtained. By subtracting the nucleus from the convex hull, a new image is created called the representative of the convex hull (ROC). Then, by Training a convolutional neural network (CNN) with the cells’ RGB image as well as the binary images of the nucleus and ROC, the generalization power is increased. The proposed method was first trained on the Raabin-WBC dataset, then its performance was evaluated on the LISC dataset without retraining. The proposed method’s accuracy on the Raabin-WBC and LISC datasets is 93.97% and 51.57 %, respectively. Besides, the generalization power of four well-known CNNs named VGG16, ResNext50, MobileNet-V2, and MnasNet1 was investigated. It was found that VGG16 has more generalization power among these models.

## Introduction

It is no secret that white blood cells (WBCs) have a central role to play in the immune system of our bodies. It can be said that WBCs are the soldiers and defenders of the body against diseases and infections. When a person gets an illness, the white blood cells come into action and try to destroy the germs and viruses. In other words, in the early stages of the disease (before appearance of the symptoms), changes in the number of WBCs in the blood are evident. Therefore, differential count of WBCs in blood samples can lead to the detection of the disease at an early stage.

In general, WBCs fall into five categories: lymphocytes, monocytes, neutrophils, eosinophils, basophils. Each of them is responsible for a special task. Lymphocytes, divided into two types of B and T [1], usually play an influential role against foreign organisms, particularly in the fight against viruses [2]. On the other hand, neutrophils serve as the primary line of defense against bacterial infections in the body [2], and eosinophils and basophils are commonly involved in inflammatory and allergic responses [2]. The typical number of WBCs in normal peripheral blood is between 4,000 to 10,000 cells per microliter, most of which are neutrophils [3]. Depending on the type of the disease, the number of WBCs either increases or decreases. Low WBC count is called leukopenia, and the high white blood cell count is called leukocytosis [4].

The analysis of normal peripheral blood samples is usually performed manually under a microscope by a hematologist. Due to the difficulty of recognizing the type of WBC, blood sample analysis is usually erroneous. Besides, the heavy workload in hematology centers and extreme fatigue have caused more erroneousness. In addition, this type of blood sample analysis is too time-consuming, and sometimes lengthening the time of blood sample analysis and examination leads to the late diagnosis of the disease. So, the disease progresses in such short time that it gets difficult to be treated. Therefore, there is a need for an intelligent, fast, and affordable system to analyze blood samples. This is where machine learning-based methods come in handy.

Usually, a proper dataset must be collected to train a classifier. After data collection, data are split into two groups of training and test. The classifier is learned with training data, and its performance is evaluated with the test set. Since the process of assembling the test set is similar to the training set, the classifier can categorize the test set accurately if appropriate and meaningful features are fed into the model. Yet unfortunately, in the real world, the classifier’s input data are generally collected with various devices or samples under different conditions than the training set. This causes a notable drop in the accuracy level of the classification.

In collecting microscopic images, blood smear staining method, camera, microscope, magnification, and lighting conditions are the most important factors that differentiate datasets from each other. These factors cause the extracted features in the two datasets to be different from each other and lead to the classifier’s making a mistake. The differences between the two datasets are sometimes so great that it is easily visible. Figure 1 shows some samples of two datasets called Raabin-WBC [5] and LISC [6], the collection processes of which are different. In Table 1, the specifications of these two datasets and their collection process have been compared.

**Table 1.** The properties of Raabin-WBC and LISC dataset [11].

| Dataset | Number of WBCs |  |  |  |  |  | Staining | Microscope | Camera |
| --- | --- | --- | --- | --- | --- | --- | --- | --- | --- |
|  | Lym. | Mono. | Neut. | Eoso. | Baso. | All |  |  |  |
| **Raabin-WBC**<br>[5] | 3461 | 795 | 8891 | 1066 | 301 | 14514 | Giemsa | 1.Olympuse CX18<br>2.Zeiss microscope (only for Basophils) | 1. Phone camera- Samsung Galaxy S5<br>2. Phone camera- LG G3 (only for basophils) |
| **LISC** [6] | 59 | 48 | 56 | 39 | 55 | 257 | Gismo-right | Axioskope40 | Sony-SSCDC50AP |

**Figure 1.**
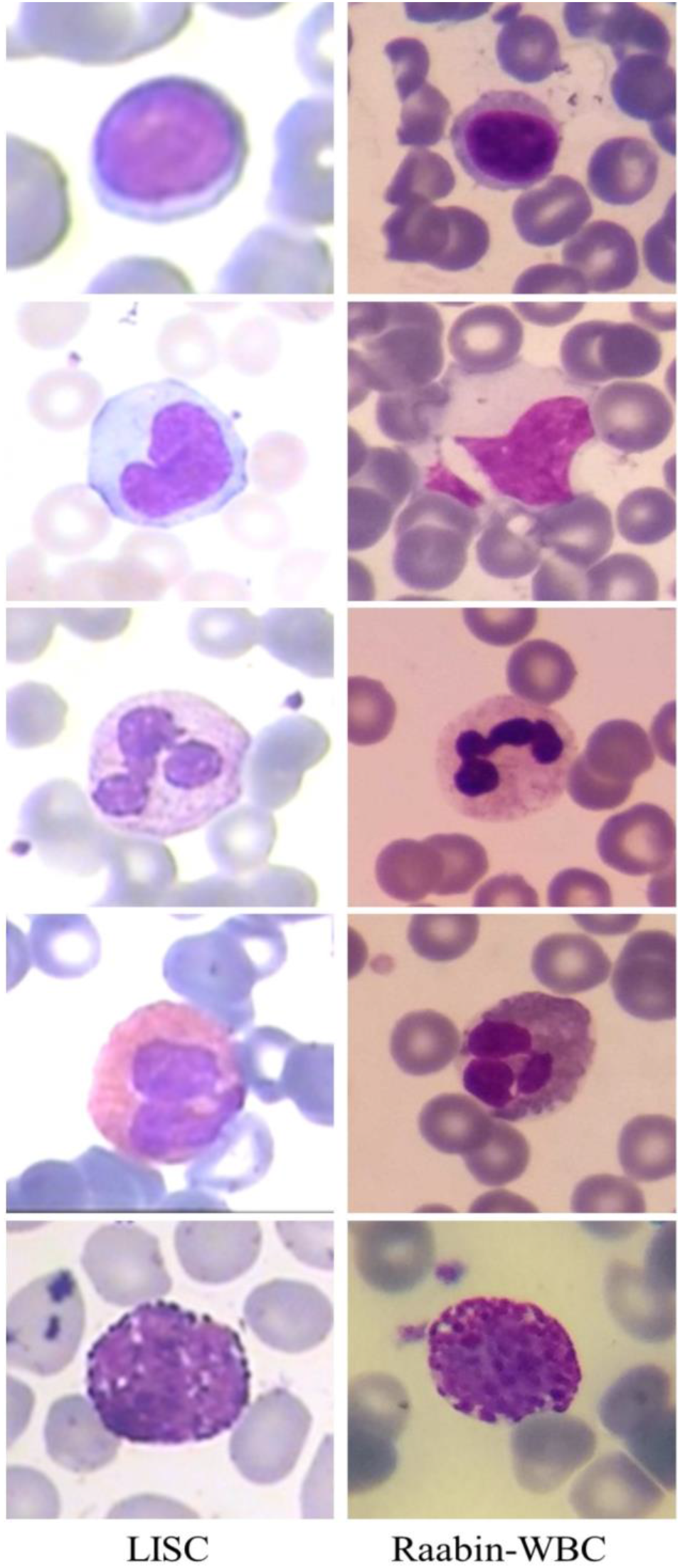
LISC and Raabin-WBC dataset. First, second, third, fourth, and fifth rows are lymphocyte, monocyte, neutrophil, eosinophil, and basophil, respectively.

Some Works have been done to overcome the low generalizability. One of these methods is transfer learning [7]. In this method, the deep neural network is trained on a very large dataset such as ImageNet [8] and then fine-tuned with a small WBC dataset. Fine-Tuning a deep convolutional neural network (CNN) trained on the ImageNet database, which contains 14 million images from 1000 categories [9], helps extract more robust and invariant and the results we reported in this article prove this claim. Domain adaptation can also be considered as an approach to increase generalizability [10]. In this attitude, the classifier still needs to be trained with a few numbers of new datasets. Another way to enhance the generalization power is to design and extract robust and invariant hand-craft features. However, developing such features is often not possible and is also a complex and time-consuming task.

Given the challenge, this paper presents a new method that has been able to classify WBCs with more generalizability. In the next section, we will elaborate on the details of this method, and then, the obtained results are presented and compared with other works.

### Research Strategy

In this study, Raabin-WBC and LISC datasets were used to investigate the generalizability of the WBC classification problem. For this purpose, training and test data of the Raabin-WBC dataset was used. Training data were employed to learn the model, and the test data were used to evaluate the classifier’s performance. Besides, to assess the generalizability of the proposed method, the LISC data were categorized using the trained classifier without further training.

### Datasets

As mentioned before, Raabin-WBC [5] and LISC dataset [6] were used for this paper. The Raabin-WBC dataset is a very large dataset with many cropped WBCs images, all of which have been labeled by two hematologists. This dataset has three sets, namely, Train, Test-A, and Test-B. We did not use Test-B set, because this set has not yet been labeled. Train and Test-A sets contain 14,514 cropped images from five aforementioned general types of WBCs. Lymphocytes, monocytes, neutrophils, and eosinophils have been collected from 56 normal peripheral blood smears. Forasmuch as basophil cells are scarce in normal peripheral blood (<1%) [1], basophil images have been collected from one cancer case (chronic myeloid leukemia). All these smears have been stained with the Giemsa technique. Also, in Table 1, the specifications of this dataset are presented with the microscope and camera type.

The LISC dataset includes 257 WBCs labeled by an expert. This dataset has been collected from peripheral blood smears and stained with Gismo-right technique [6]. These images have been imaged at a magnification of 100 employing a light microscope (Microscope-Axioskope 40) and a digital camera (Sony Model No. SSCDC50AP) [6]. The properties of the LISC dataset have been presented in Table 1.

### Steps of the proposed method

The proposed method contains three steps as following:

- Detecting and segmenting the nucleus
- Obtaining the convex hull of the nucleus and subtracting the segmented nucleus from the convex features and adds more generalization power. Although this approach has had some successes, it is still not satisfying, hull for making a new binary image. We named this image as the representative of the convex hull (ROC).
- Classifying WBCs through feeding the RGB images, segmented images of the nucleus and ROC images into a CNN.

We will elaborate more on the proposed method in the next sections.

### The nucleus segmentation algorithm

To segment the nucleus, we utilized the method used in our previous work [12]. In this method, firstly, the RGB image is converted to CMYK color space. After that, two threshold values are computed using Otsu’s thresholding algorithm [13]. The first threshold is obtained by applying two-class Otsu’s thresholding algorithm, and the second threshold is obtained employing three-class Otsu’s thresholding algorithm. Finally, the ultimate threshold value is attained through calculating the combination convex of the first threshold value and the second threshold value. This method can detect the nucleus with a dice similarity coefficient of 95.42 %. More details of the segmentation algorithm have been presented in [12].

### Obtaining ROC image

By subtracting the segmented nucleus from its convex hull, the ROC image is obtained. However, here lies a big question why we made and used this image. It is clear that the shape of the nucleus is an important feature to diagnose the type of WBC. Considering this hypothesis, we found that the ROC image can also help to classify WBCs better. Figure 2 illustrates this claim. We can see that the ROC for the lymphocyte is very different from the ROC for the neutrophil.

**Figure 2.**
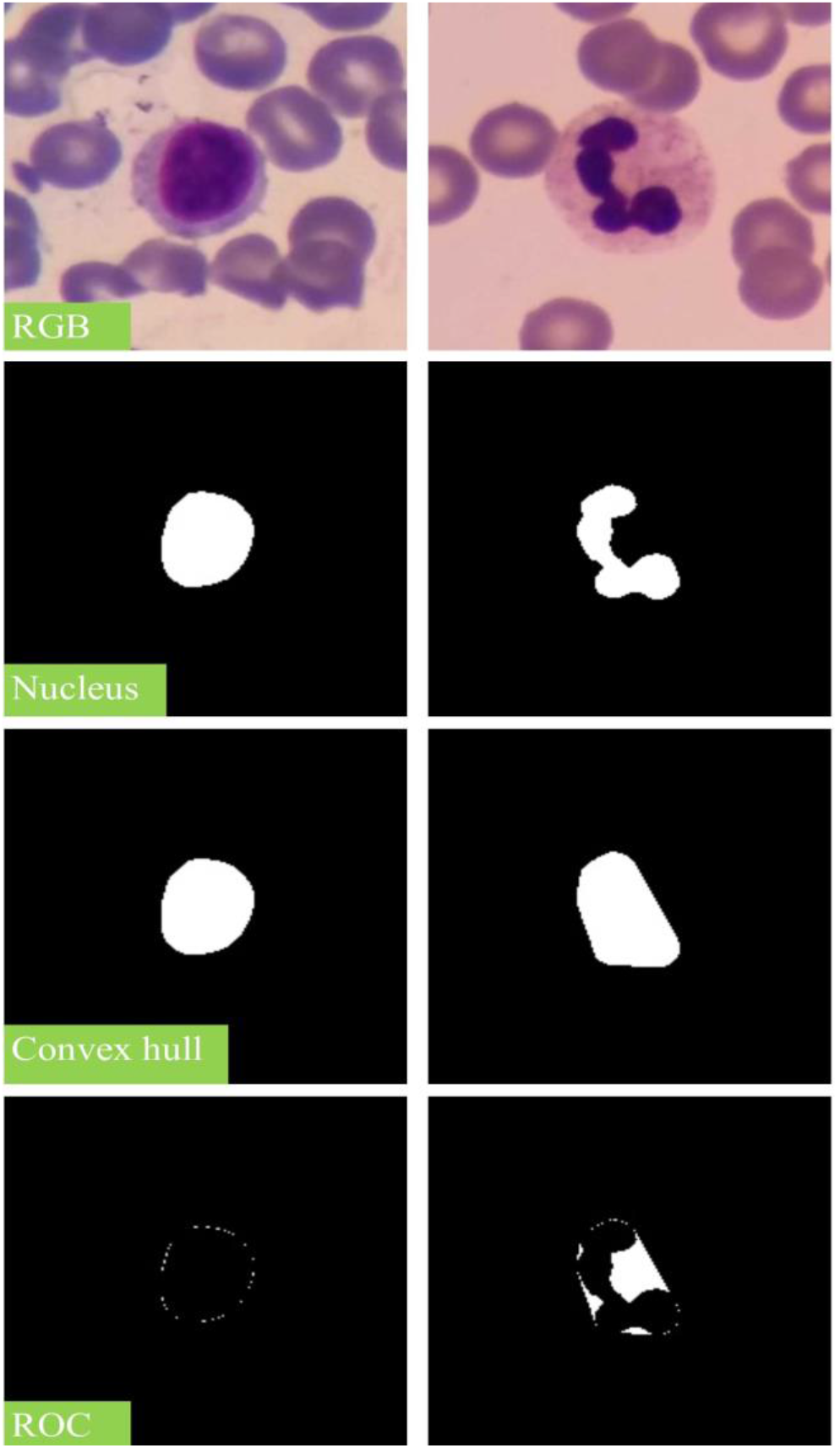
The segmented nucleus (first row), the convex hull of the nucleus (second row), and the ROC image

### Classification

To classify WBCs, we used convolutional neural network. Three channels of RGB image are usually inputted to the CNN, but in this paper, beside the RGB channels, we injected the binary images of the nucleus and ROC to the model. Therefore, this CNN model was created so that its input shape was 200×200×5. This network contains seven convolutional and three fully connected layers. Furthermore, to prevent over fitting, three batch normalization layers were employed. The training process was performed through stochastic gradient descent algorithm with a momentum coefficient of 0.9. This network has more than 2.2 million trainable parameters. Figure 3 shows the structure of the model.

**Figure 3.**
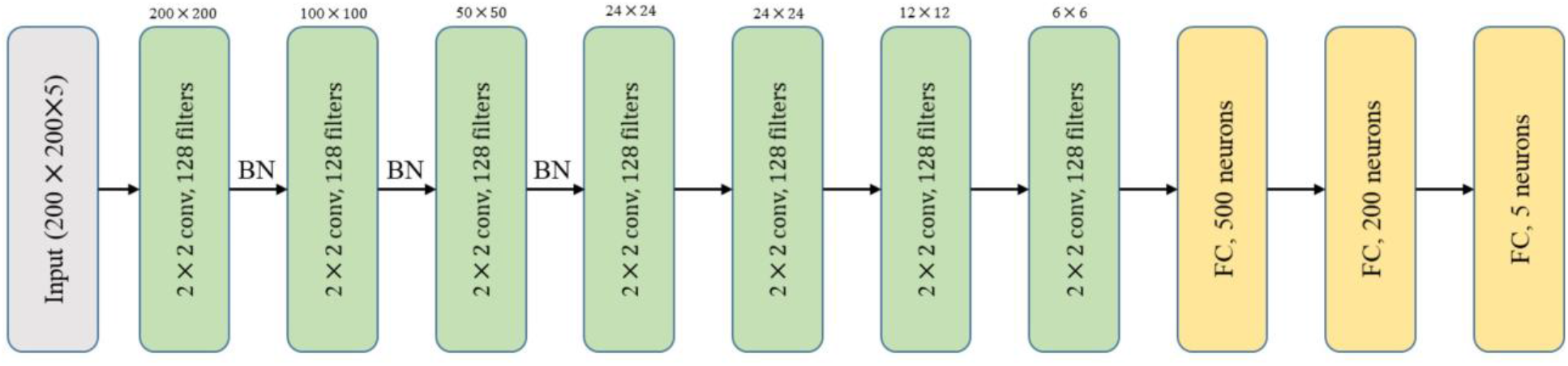
Architecture of the CNN model used in this paper. Conv (convolutional layer). BN (batch normalization layer). FC (fully connected).

The Raabin-WBC dataset was already divided into two parts for learning the model: about 70 percent for the training set and about 30 percent for the test (Test-A set). Moreover, 15 percent of the training set was considered as a validation set during learning process. The CNN model was trained for 15 epochs.

The Raabin-WBC dataset was used to train the model and evaluate its performance, and the LISC dataset was utilized to examine the proposed method’s generalizability. It is worth noting that the model was not retrained with the LISC dataset.

The accuracy curves have been plotted for each epoch (Figure 4 and Figure 5). From Figure 4 and 5, it can be seen that there is a dramatic drop in accuracy for the LISC dataset.

**Figure 4.**
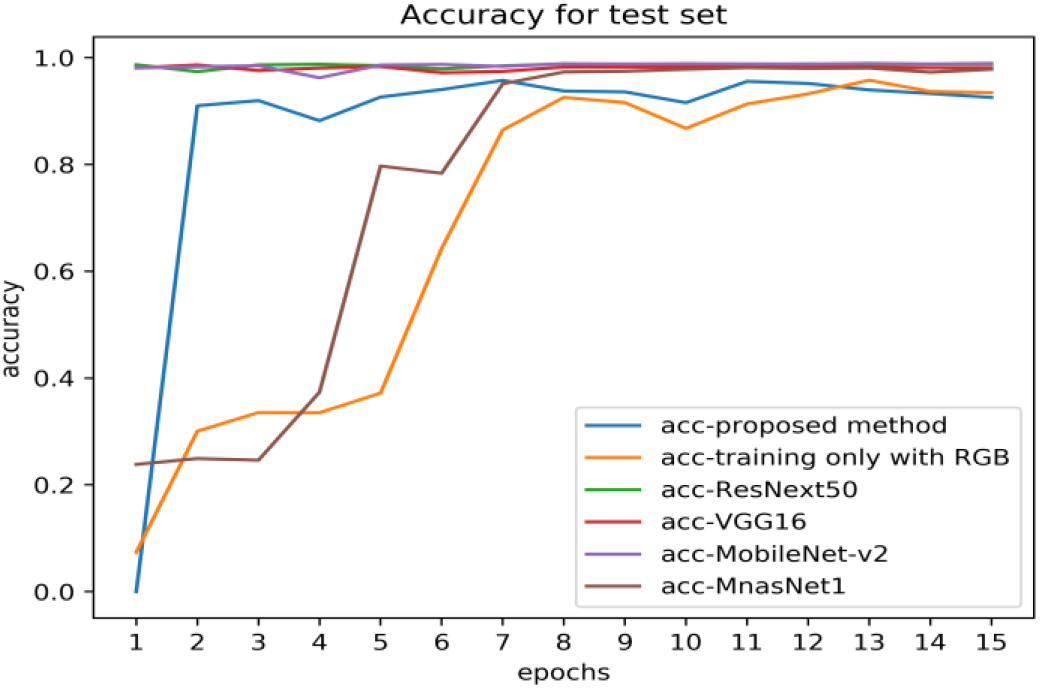
The accuracy curve for test set

**Figure 5.**
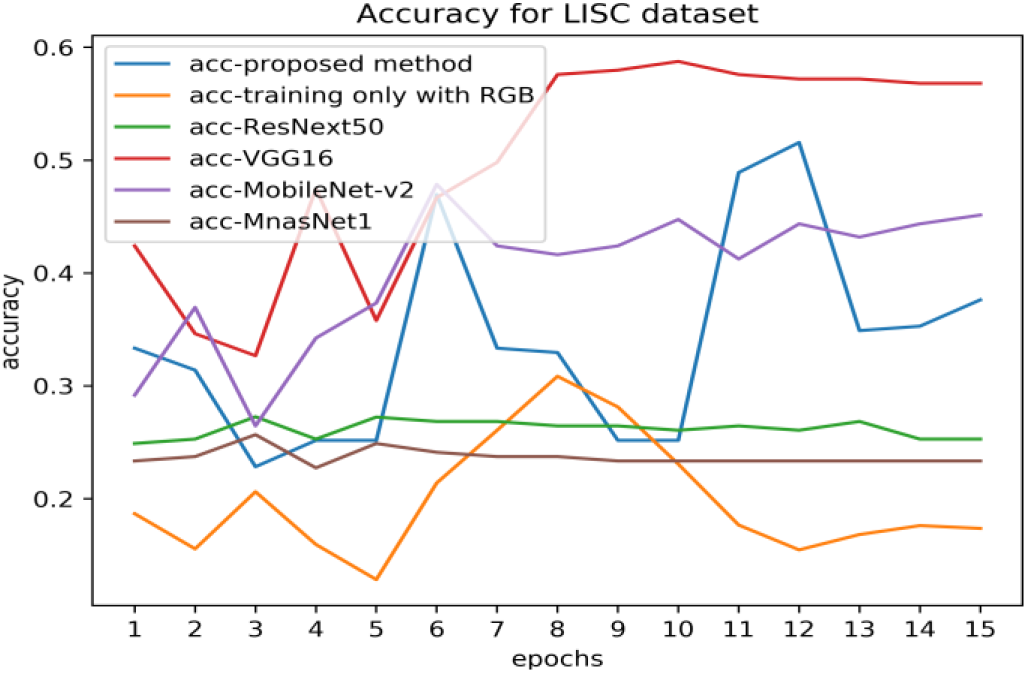
The accuracy curve for LISC dataset

## Results and Discussion

In order to have a proper comparison, we trained the convolutional network designed for this study with RGB image channels, only, and examined the generalizability in this case. Moreover, we investigated the generalizability of transfer learning-based methods employing four well-known deep CNNs called VGG16 [14], ResNext50 [15], MnasNet1 [16], and MobileNet-v2 [17]. For this purpose, these networks were fine-tuned with training data. These models have already been trained on the ImageNet dataset, which contains 14 million images from 1,000 classes. These networks are expected to have high generalizability because they have seen many different images, but the results show otherwise.

From Table 2, Figure 4 and Figure 5, it can be seen that the accuracy of all models has dropped dramatically. By taking a meticulous look at Table 2, we can see that the accuracy of the model trained with RGB image fell from 86.76 % to 30.85 %, while the accuracy of the proposed method dropped from 93.97 % to 51.57 %.; it reveals that our approach towards injecting the binary images of the nucleus and ROC has been helpful in increasing the generalizability. In addition, among all the pre-trained networks, VGG16 had the best generalizability. As can be seen, by changing the dataset, the accuracy of this network, albeit having the best generalization power among all models, decreases from 99.91 % to 58.75 %.

**Table 2.**
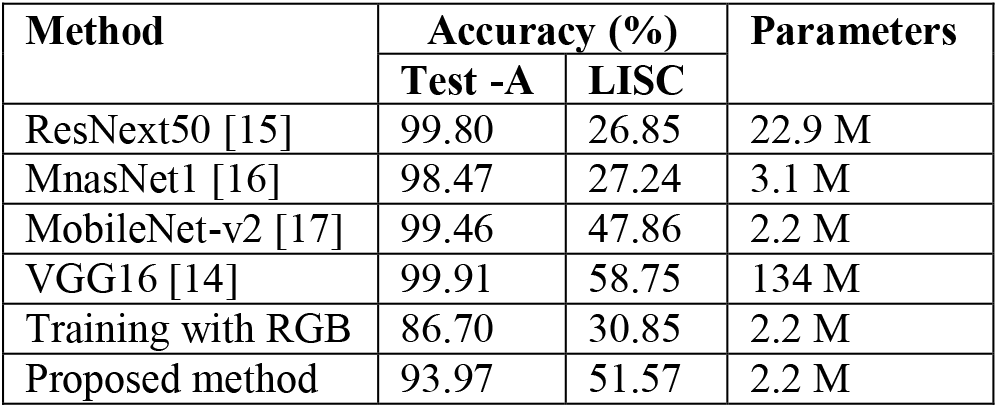
The comparison of generalization power. M (million)

Our proposed method ranked second among all these models after the VGG in terms of generalizability. It should be noted that the network used in our method has only 2.2 million trainable parameters. In comparison, the VGG16 network has 134 million trainable parameters, and this network was also trained on the ImageNet database. Therefore, holding the second position is not bad and shows that the proposed method has been successful in performance.

Injecting binary images of the nucleus and ROC to the CNN led to an increase in the generalizability; it could be because the shape of the nucleus does not change with changing the dataset. The binary image is also resistant to changing the dataset, because its pixels are either 0 or 1. Therefore, the camera, microscope, staining method, and lighting conditions can not affect the binary image of WBC’s nucleus. As a result, this approach is useful and has helped generalization power to improve.

It is worth noting that all the results reported in Table 2 are considered as the best accuracy level of each model for the LISC dataset (best epoch).

## Conclusions

This paper attempts to provide a new efficient method to increase generalizability. The results showed that our proposed method (using binary images of the nucleus and ROC) is effective in increasing generalization power. Moreover, we investigated the generalizability of four prominent CNN models and illustrated that the VGG16 has more generalizability than other models reported in Table 2.

### Implementing environment

Our method and all the pre-trained networks mentioned in Table 2, were implemented using Python 3.6.9 and Pytorch library version 1.5.1.

